# epiTAD: a web application for visualizing high throughput chromosome conformation capture data in the context of genetic epidemiology

**DOI:** 10.1101/243840

**Authors:** Jordan H. Creed, Garrick Aden-Buie, Alvaro N. Monteiro, Travis A. Gerke

## Abstract

The increasing availability of public data resources coupled with advancements in genomic technology has created greater opportunities for researchers to examine the genome on a large and complex scale. To meet the need for integrative genome wide exploration, we present epiTAD. This web-based tool enables researchers to compare genomic structures and annotations across multiple databases and platforms in an interactive manner in order to facilitate *in silico* discovery. epiTAD can be accessed at https://apps.gerkelab.com/epiTAD/.

## 1 Introduction

Epidemiological studies have provided strong evidence for a genetic component in the predisposition to common diseases. Recently, Genome-wide Associations Studies (GWAS) have identified a large number of chromosomal loci defined by Single Nucleotide Polymorphisms (SNPs) (https://www.ebi.ac.uk/gwas/)[1]. Although several thousands of these SNPs have been associated with different traits at genome-wide statistical significance (5 x 10^-8^), the mechanistic underpinnings of these associations are still largely unexplored.

The picture emerging from representative functional analysis of GWAS SNPs is the one in which these SNP are enriched in genomic regions with evidence for transcriptional regulatory activity, such enhancers and promoters. In this scenario, trait-associated SNPs exert their allele-specific effects through transcriptional regulation of (mostly unknown) target genes by modifying transcription factor binding sites, in turn changing the activity of the enhancer/promoter region region[2, 3, 4]. Based on these conclusions, it becomes clear that reliable identification of enhancer target genes will be instrumental for functional dissection of GWAS loci[2].

The comprehensive identification of regulatory regions active in multiple cell lines conducted by ENCODE[5, 6, 7] and the GTEX project[8] have accelerated analysis of these loci by providing large resources of putative enhancers and potential target genes. Linking an enhancer region to a candidate gene has been performed using two main approaches: expression quantitative trait loci (eQTL) analysis, which is based on a statistical association between genotypes and expression levels; and biochemical techniques, such as chromosome conformation capture[9] (3C and related techniques) or Chromatin Interaction Analysis by Paired-End Tag Sequencing (ChIA-PET).

However, expression data from GTEx is obtained from a whole tissue and thus reliable expression estimates for cells with low representation in the tissue (e.g. melanocytes in skin) are not available. Moreover, because of the dynamic nature of gene expression, some enhancers may only be active in certain conditions or restricted developmental windows

Recent studies of interphase human chromosomes has revealed a hierarchical organization with the genome largely being divided into sections of open and closed chromatin[10], which are then organized by Topologically Associating Domain (or TADs), and can be further organized as insulated neighborhoods or loops. TADs have been shown to be mostly independent of cell type and, importantly, cis-regulatory effects mediated by chromatin looping interactions between enhancers and promoter regions tend to happen within topologically associated domains[11].

Thus, determination of which TAD includes the trait-associated SNPs also has the ability to narrow down the list of candidate genes. Further, as the resolution of HiC increases, investigators should also be able to identify specific interactions between enhancers and promoters. However, until now there was no available tool to integrate data of epidemiologic interest (e.g. GWAS SNps, eQTL, Haploreg, and RegulomeDB) with HiC data. The tool described here aims to integrate these different data sources.

## 2 Materials and Methods

### 2.1 Data Collection

The Hi-C reads provided in epiTAD are from human fibroblast IMR90 cell profiles as described by Dixon et al[11]. TAD boundaries were also taken from the Dixon experiments and were determined on the basis of a “bi-directionality index” of chromatin interactions[11]. Linkage disequilibrium (LD) measures are queried from HaploReg, with data originating from the 1000 Genomes project[12]. eQTLs are obtained through GTEx Version 7, which determines eQTLs from 635 samples over 53 tissues[13]. Coordinates and HGNC symbols for genes are taken from ENSEMBL, while data from various sources are scraped from the annotation aggregator Oncotator. All data is queried in real time from publicly available resources.

### 2.2 Implementation

epiTAD is available for public use online at the author’s website https://apps.gerkelab.com/epiTAD, and a containerized version of the application is archived and available via Docker Hub https://github.com/GerkeLab/epiTAD/Dock The user interface is built using R Shiny [14], and real-time data querying, table production and visualization is enabled by the R packages haploR [15], HiTC [16], Sushi [17], and biomaRt [18]. Full source code for the application is available at https://github.com/GerkeLab/epiTAD.

## 3 Access and Display

epiTAD is freely available online from https://apps.gerkelab.com/epiTAD/, and is compatible with all major web browsers (Chrome, Internet Explorer and Safari). No login or user information is required for use. The display consists of two pages: the primary page, which contains all tables, visuals, and user inputs; and an informational page, accessible from a tab on the left-hand side of the page. The tool is broken down into five panels: inputs, output options, variant annotations, gene annotations, and visualization.

At a minimum, users need to input at least one SNP, in the form of a dbSNP rs id (e.g. rs10486567). Multiple SNPs can be uploaded and queried at once either as a comma separated list or as a text file, however they must all reside on the same chromosome.

Variant annotations from the HaploReg and RegulomeDB databases are available under separate tabs. The amount of information shown can be customized by the user from the “Select Output” panel under the tab of the same name. The TADs tab informs the user if the SNP(s) are located within a known TAD and if so, provides the coordinates and also contains a link to the Hi-C Browser for additional visualization options.

A query region is created that contains all SNPs within the maximum of the LD or TAD boundaries. If there are no SNPs above the selected LD threshold and the variants of interest are not located within a TAD, then a region of 53,500 bp is added to either side of the SNP to create the query region. This query region is then used for all gene level annotations and for the initial coordinates for the visualization. The gene names and coordinates can be downloaded from the site as a CSV file. Oncotator queries are broken down by source and can pull information from the Cancer Gene Census, HUGO Gene Nomenclature Committee, and UniProt. If eQTLs are available from GTEx the user can select tissue(s) of interest to subset results.

Live links to additional resources and tools, including ClinVar, the UCSC Genome Browser, Juicebox, Hi-C Browser, and GTEx, are made available to the user for the region/SNPs.

A figure is automatically rendered that spans the query region and contains four tracks. The first track displays the Hi-C contact matrix, the second contains gene coordinates and names within the region, the third shows the query SNP(s) (color) and SNPs in LD (grey), and the final track shows the TAD boundaries. The plot can be updated with user-defined coordinates, as long as the coordinates are at least 20,000 bp apart (Hi-C is binned in 10,000 bp), and there is a button to return to the original coordinates. The user also has the option to choose from 5 color schemes. The figure can also be downloaded as a high-resolution PDF.

Users can bookmark any querys to return to the page setup as-is, saving all edits or selections made. Figure 1 demonstrates the output for rs10486567. This SNP was previously found to significantly associate with *HIBADH* expression in prostate tumor tissue and with *TAX1BP1* expression in normal prostate tissue[19]. This finding may be, in part, explained by observing that both of these genes as well as the query SNP are located within the same TAD, with the Hi-C data showing a large amount of contact across this area.

**Figure 1:**
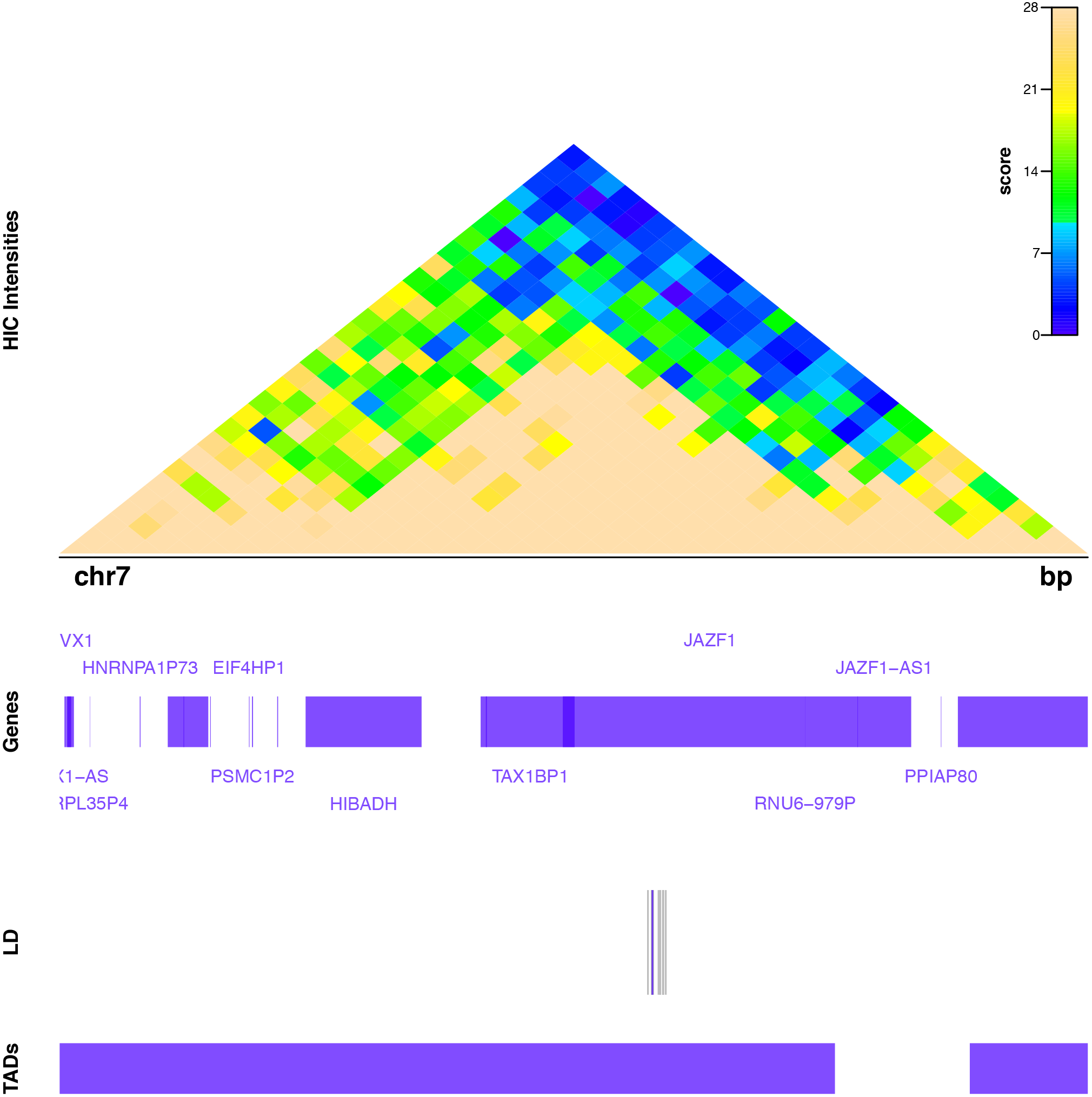
An example of the epiTAD _gure output for rs10486567.

## 4 Comparison to other tools

Various web tools and stand-alone programs enable the visualization of a researcher’s Hi-C data, or utilize competing algorithms to call TADs; however, they do not adequately show the relationship between these measures and other units of genomic organization. The Hi-C Browser[20] and Juicebox[21] are other free web tools for visualizing Hi-C data. These tools and epiTAD produce heatmaps, but differ in additional annotations and data sources provided. Juicebox is strongest when used as either part of Aiden Lab’s Hi-C pipeline or as a stand alone program for visualizing published experimental data in a single image, while Hi-C Browser contains a unique track for DNase I hypersensitive sites (DHSs). Both tools also allow users to either select specific cell lines or use their own data. While these tools are highly valuable, they may be of limited use to researchers with an interest in variant level genome structure or population genetics, as neither tool delivers such annotations. Furthermore, high-resolution figures are not downloadable. epiTAD serves to fill this role for researchers.

## 5 Discussion

In light of the ongoing and rapid growth in publicly available genomic data, efficient systems to compare genomic databases become increasingly vital to researchers. epiTAD provides a single application to integrate genomic annotations and measurements across multiple public databases covering a uniform region. The application allows researchers to access and plot large amounts of data related to major genome organization structures without programming knowledge. The resulting web app may prove a broadly useful component of in *silico* functional genomics discovery.

